# EPIFANY – A method for efficient high-confidence protein inference

**DOI:** 10.1101/734327

**Authors:** Julianus Pfeuffer, Timo Sachsenberg, Tjeerd M. H. Dijkstra, Oliver Serang, Knut Reinert, Oliver Kohlbacher

## Abstract

Accurate protein inference under the presence of shared peptides is still one of the key problems in bottom-up proteomics. Most protein inference tools employing simple heuristic inference strategies are efficient, but exhibit reduced accuracy. More advanced probabilistic methods often exhibit better inference quality but tend to be too slow for large data sets.

Here we present a novel protein inference method, EPIFANY, combining a loopy belief propagation algorithm with convolution trees for efficient processing of Bayesian networks. We demonstrate that EPIFANY combines the reliable protein inference of Bayesian methods with significantly shorter runtimes. On the 2016 iPRG protein inference benchmark data EPIFANY is the only tested method which finds all true-positive proteins at a 5% protein FDR without strict pre-filtering on PSM level, yielding an increase in identification performance (+10% in the number of true positives and +35% in partial AUC) compared to previous approaches. Even very large data sets with hundreds of thousands of spectra (which are intractable with other Bayesian and some non-Bayesian tools) can be processed with EPIFANY within minutes. The increased inference quality including shared peptides results in better protein inference results and thus increased robustness of the biological hypotheses generated.

EPIFANY is available as open-source software for all major platforms at https://OpenMS.de/epifany.

## Introduction

Ever since the emergence of bottom-up proteomics experiments^1^, mapping the identified peptides back to their most plausible source proteins, the protein inference problem, has been a key problem in proteomics^2–4^. High dynamic range of protein abundance, limitations in digestion, separation, and mass spectrometry result in incomplete coverage of the source proteins by identified peptides. Reconstructing the source proteins originally present in the sample should thus rely on as much of the experimental evidence (i.e., peptide identifications) as possible – which also includes non-unique peptides shared between multiple source proteins. Starting from a notion of a probability of the presence or absence of peptides in the sample, usually expressed by a score, we want to infer the presence or absence of the proteins these peptides originated from. Due to the common presence of ambiguous peptides arising from one or more proteins sharing parts of their amino acid sequence this is not a trivial task. ^2^

The scores for peptides are typically obtained by so-called peptide search engines that match experimentally observed spectra to theoretically derived ones based on the sequences of an in silico digested database of protein candidates. Those peptide-spectrum matches (PSMs) then need to be scored to be able to quantify the uncertainty in correctness of such a match. Uncertainty in the assignment of a peptide sequence to a spectrum may be a consequence of multiple peptide candidates matching to the same spectrum or a result of imperfect data such as incomplete or noisy spectra as well as incomplete protein databases. ^5^

The formulation of protein inference algorithms naturally leads to a representation of the relation between peptides and proteins as a bipartite graph of nodes (proteins and peptides) that are connected with an edge if a peptide is part of the theoretical digest of the (parent) protein (Figure 1). Figure 1 also shows that the ambiguity of peptides across proteins may lead to proteins without unique evidence (e.g., protein E) and in an extreme case to experimentally indistinguishable protein groups (e.g., one comprising proteins F and G, which share all their observed peptides). Reasons for ambiguous peptides are manifold in biology and include among others homology, alternative splicing or somatic recombination. Depending on the degree of ambiguity between the peptides of different proteins they are often clustered into various types of so called protein ambiguity groups.^2^ It should be noted that those groups can either be defined solely based on the experimentally observed peptides or based on all theoretically possible peptides.^6^ Preliminary grouping (especially of indistinguishable proteins) is often used automatically by inference algorithms to solve a less ambiguous problem. While this is enough for studies that only need to confirm presence of any protein in a group (using group level inference probabilities), other studies look for the effects of a certain isoform on a disease and need to know how likely it is that this specific isoform was expressed (therefore interested in single protein level results). To compare results of inference methods on the same level (wherever possible) is an important consideration during benchmarking.

**Figure 1:**
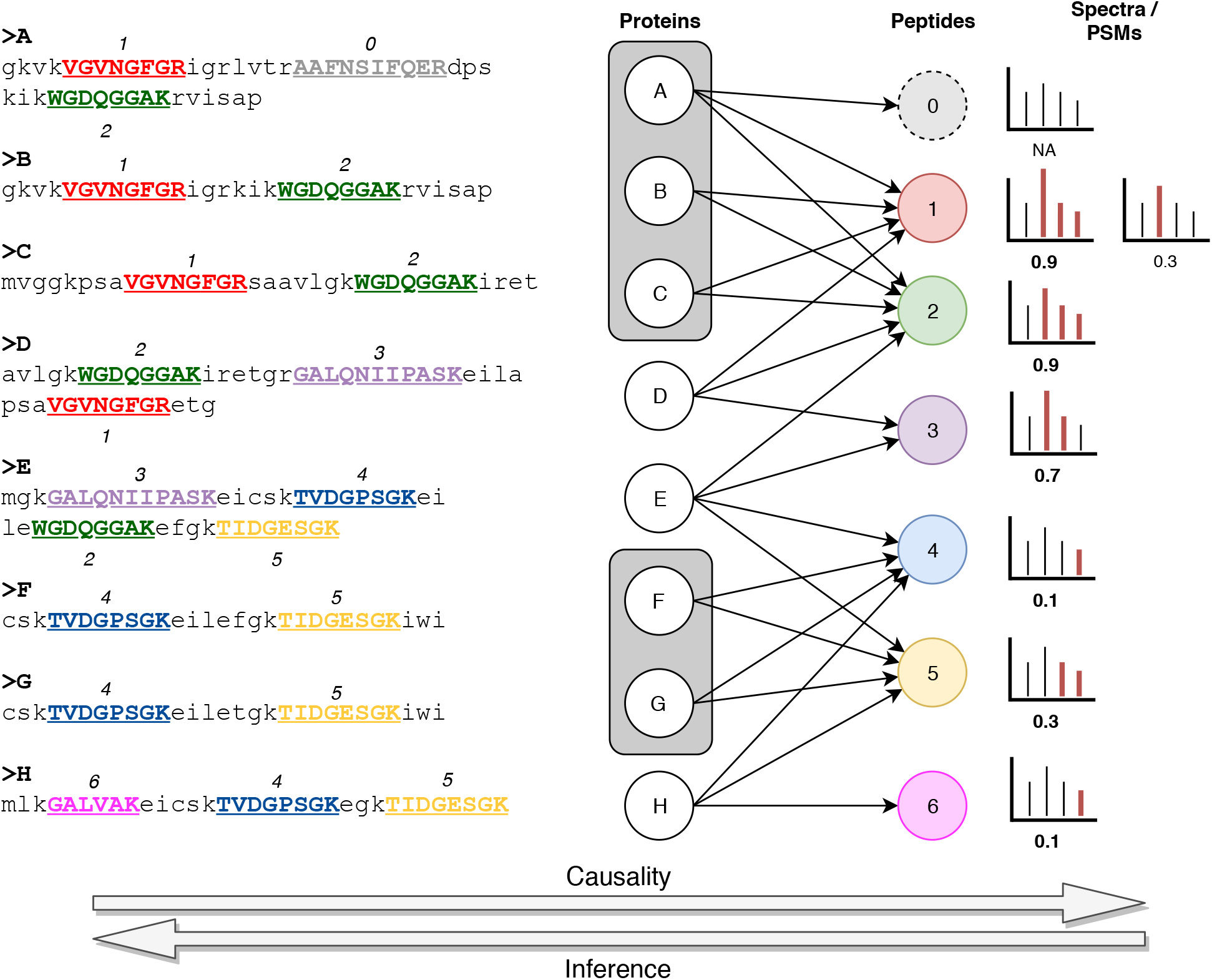
Example of a bipartite protein-peptide graph. Nodes with letters represent potential proteins from the input database. Colored nodes are the peptides from in-silico digest with the given enzyme (trypsin). Arrows are drawn when a protein may theoretically generate a peptide. Dashed circles represent experimentally unobserved entities due to missing (NA) peptide-spectrum matches. Red peaks in the sketched hypothetical tandem mass spectra were matched to a theoretical ion of the peptide that matched best to this spectrum. Probability scores roughly follow a dot-product based score but were invented for the sake of this example. Bold scores highlight the chosen match probability for this peptide (i.e., the maximum probability). The left side shows the used protein database with their tryptic peptides (upper-case bold underlined substrings) following the same color and number scheme as the nodes in the graph. Proteins in the same shaded curved rectangle comprise an experimentally indistinguishable (ambiguity) group. The arrows on the bottom show the general directions of the two processes: causality in the course of the experiment and inference based on the observed data.

Early inference approaches resorted to simple rule-based conclusions. If a protein is connected to *n* or more peptides—where *n* is usually one or sometimes two to avoid so-called one-hit-wonders—then it is declared present, otherwise absent. The problem with such approaches, however, is the implicit overcounting of shared peptides. In the presence of very large proteins like titin, false positive identifications may arise due to matching one of its many peptides. Similarly, titin might be wrongly identified if it just shares enough peptides with truly present proteins. Some methods tackle this problem by either ignoring shared peptides (Percolator^7,8^), employing maximum parsimony principles and finding a minimal set of proteins explaining found peptides or PSMs (PIA^4^), iteratively distributing its evidence among all parents (ProteinProphet^9^) or incorporating the evidence in a fully probabilistic manner (Fido^10^, MSBayesPro^11^, MIPGEM^12^) to make use of the “explaining-away” property of Bayesian networks^13^. “Explaining-away” is a term used in probabilistic reasoning to describe the implicit conditional dependency between multiple causes of a common effect (when its probability is non-zero). In this case knowledge about one cause from other evidence or prior to inference influences our belief about the other causes. In probabilistic models with synergistic parametrizations like the ones in the aforementioned Bayesian tools this means that if one protein is very likely to be present from its unique evidence, it is already a sufficient explanation for peptides that it shares with other proteins (without evidence) and thereby affects their probability in a negative way. This leads to a probabilistic type of parsimony.

On a gold standard dataset it was shown by The et al. ^14^ that fully probabilistic models perform among the best in terms of the pure identification task. However, current solutions are computationally demanding. Additionally, in the case of candidate protein databases with many cases of peptides being shared between truly present and absent proteins, the reported probabilities are not a good basis for well-calibrated target-decoy false discovery rates (FDRs) as they yield poor approximations of the true FDR. This leads to over-/underestimation of the true amount of false discoveries.

In our new approach EPIFANY (Efficient Protein InFerence for ANY protein-peptide network) we used a fast approximate inference algorithm called loopy belief propagation (LBP) which has already been shown to perform well on solving other types of probabilistic graphical models (e.g., models used in important information theoretic algorithms like the error-correcting turbo codes^15^ as well as on quick medical reference (QMR) disease diagnosis networks^16^). Using LBP we can achieve drastically improved runtimes than other Bayesian approaches without any approximations on the underlying graph itself. We improved the calibration of the resulting FDRs by introducing an optional regularized model with max-product inference and a greedy protein group resolution based on the reported protein probabilities.

## Methods

In the following section the underlying probabilistic graphical model, the inference procedure as well as the pre- and post-processing steps on the data are explained in detail.

### Model

The model we chose for protein inference is based on a Bayesian network (BN) representation. The protein-peptide graph (Figure 1) encodes the conditional dependencies of proteins and their peptides. The advantages of this specification of conditional (in-)dependencies is the resulting factorization of the high-dimensional joint distribution into smaller distributions, namely prior distributions for the proteins and conditional probability distributions (CPDs) for the peptides given their parents. In case of the binary representation for every peptide and protein, these distributions are discrete and correspond to probability tables.

Additionally, the Bayesian network needs to be parametrized. Although the factorization into smaller distributions decreases the number of parameters, each CPD still needs 2^*p*^ parameters, where *p* is the number of parent nodes, to be set or learned. By recognizing the fact that in the generative process from proteins to peptides the presence of any of the parent proteins is enough to potentially produce a peptide, we can reduce the number of parameters further when specifying the conditional probability according to a noisy-OR model^13^. In its original form, the network using the noisy-OR model requires the following parameters:

- *γ*_*ρ*_ prior for protein with index *ρ*
- *α*_*ρ,ε*_ noisy-OR emission probability of a protein *ρ* generating peptide *ε*
- *β*_*ε*_ noisy-OR leak probability for a peptide *ε* being generated by chance

For now, we employ the same simplification used in the Fido^10^ algorithm by assuming equality among all *α* and the presence of only one *β*. Additionally, constant priors *γ* for all proteins prevents biases on protein level when no further information is available. The parameters *α, β, γ* either have to be specified manually, or are by default selected from a grid of initial values based on target-decoy classification performance and probability calibration (see Implementation subsection for details).

With these assumptions, the noisy-OR model suggests Equations 2 and 3 below for the CPD of a specific peptide *ε* given the presence of its parent proteins. The binary random variable *E*_*ε*_ denotes the presence of peptide *ε* while the binary random variables *R*_*ε*,1_,…, *R*_*ε,p*_ represent the presence of the parent proteins of peptide *ε*. *N*_*ε*_ is a random variable for the total number of present parent proteins of a peptide.

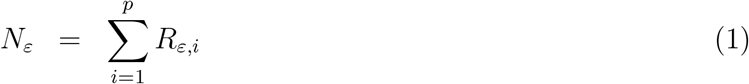

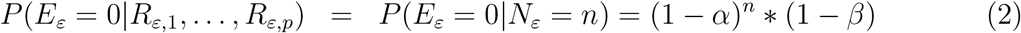

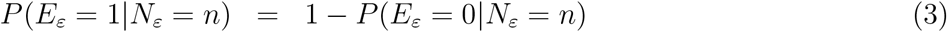

Note that Equation 2 is exploiting the symmetry arising due to an equal *α* for all proteins of a peptide. One addition to the original model is an option to put regularizing priors of the following unnormalized form onto the number of proteins that may produce a certain peptide *ε*:

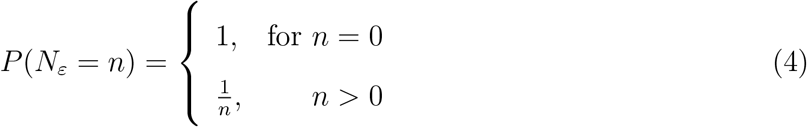

This results in a more uneven distribution of the evidence from peptide *ε* starting at the most likely producing proteins (based on their beliefs from the rest of the network), especially in conjunction with max-product inference.

### Algorithm

The goal of creating a representation of the problem as a (factorized) probability distribution is to eventually perform inference on random variables of interest. In this specific case, the probabilities of interest are the marginal probabilities of the proteins given the evidence on the peptide level, the so called posterior probabilities.

As peptide level evidence, the algorithm first reads the PSM probabilities and the associations to their parent proteins from spectra searched with a peptide search engine. It then aggregates PSM probabilities on the peptide level by picking the maximum PSM probability per unmodified peptide sequence and filters out peptides with extremely small probabilities (e.g., below 0.001).

A naive approach to inference in a Bayesian network would be to create a large joint probability table for each connected component of the network and marginalize for each protein by aggregating the probabilities of all possible configurations leading to the same state of the current protein of interest. To make this approach viable for at least small to medium-sized problems, previous tools resorted to (Gibbs) sampling ^11^ or sped up calculations by caching results and making use of symmetries arising due to the chosen model^10^. A possible symmetry to be exploited is the dependence of the peptide probability only on the number of parent proteins (not their exact combination) as can be seen in Equation 2. This symmetry consideration reduces the number of different input configurations in the presence of indistinguishable groups. Although this procedure has been implemented efficiently in tools like Fido, the worst-case runtime is exponential in the number of proteins in a connected component. Reducing the size of the connected components by splitting them at low-probability peptides is a reasonable approximation only if the probability cutoff is not too high. Using the looping version^16^ of Pearl’s belief propagation algorithm^17^, even non-tree structured graphs with cycles (such as all but the most simple protein-peptide networks) can be processed efficiently while keeping flexibility in which types of factors (probability table-based factors, function-like factors, convolution-tree-based adder factors, etc.) on which sets of random variables are used. Convolution trees^18^ (CTs) are an important means to efficiently calculate the sum of discrete probability distributions (e.g., to apply Equation 1). The general idea of applying loopy belief propagation to our problem is to create a factor graph (see Figure 2 and Implementation subsection) out of the specified Bayesian network and initialize messages on all edges in both directions uniformly. All so-called factor potentials are initialized according to their priors and/or likelihoods. Lastly, the algorithm iteratively queues, updates and passes messages between the factors to be incorporated into their potentials until messages do not change anymore (i.e., convergence is reached in terms of their mean-squared error). By default, messages with the highest residuals gain highest priority for the next iteration^19^. Potentials *ψ* are updated by incoming messages *φ* following the HUGIN algorithm^20^ according to the equation below for a message from node with random variables (RVs) *V* to a node with RVs *W* whose intersection is *S*:

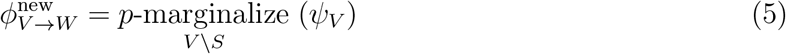

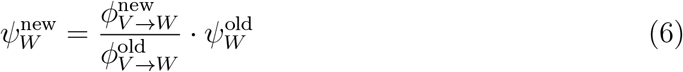

*p*-marginalization is a generalized form of marginalization where instead of summing over removed variables (equivalent to *p* = 1) the *p*-norm is computed. To reach convergence faster, messages can be dampened (by a “momentum”) in a way that the updated message is a convex combination of old and new messages^16^.

**Figure 2:**
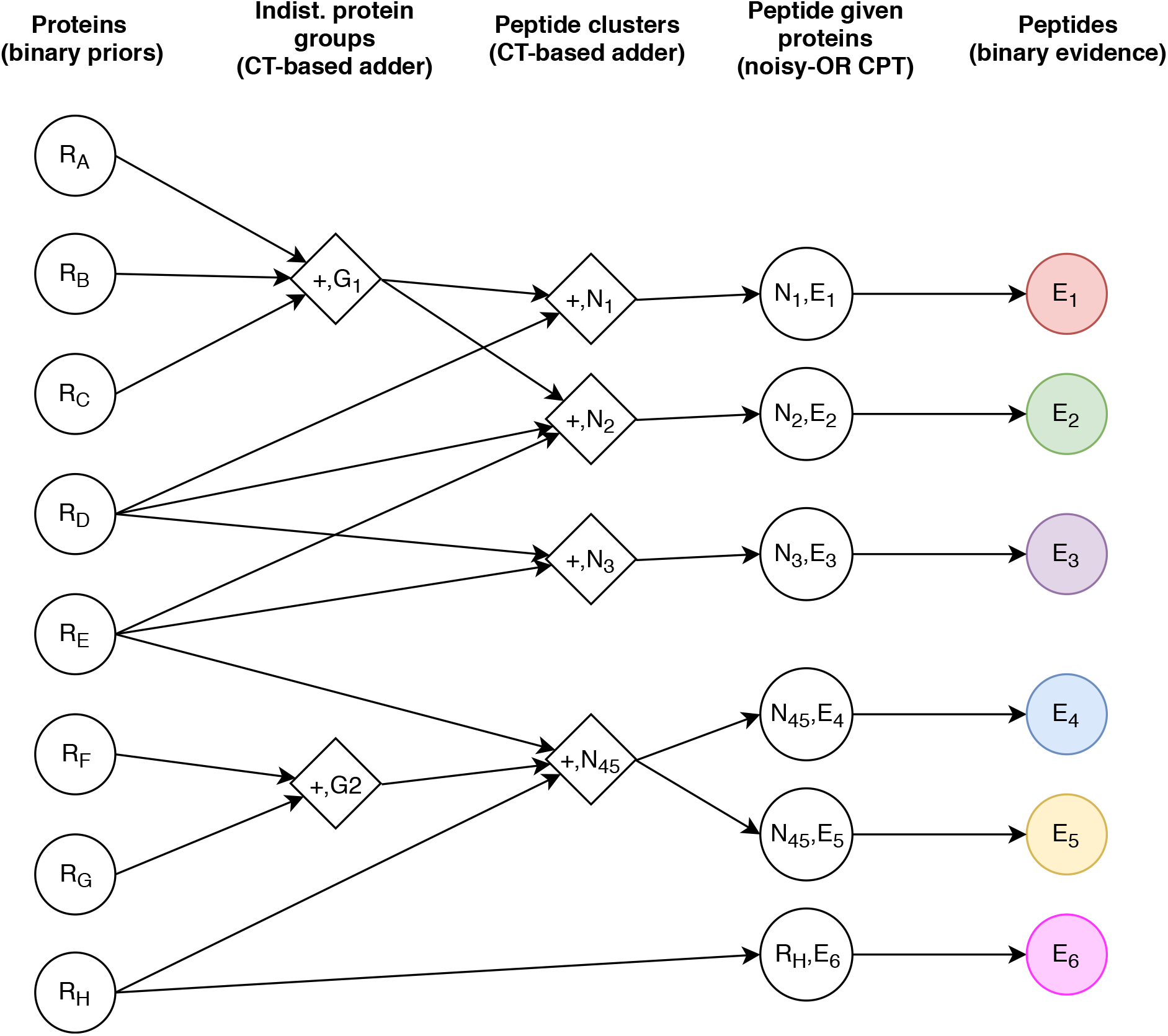
The factor graph created for the loopy belief propagation algorithm based on the example in Figure 1 (same color and letters). Each node represents one factor. Annotations on the factors describe the set of random variables (RVs) comprising its potential. Circles are table-based factors, diamonds convolution-tree (CT) based probabilistic adders. The set of RVs on CT-based adders are implicitly defined by the union of variables from neighbors on the left side of the graph (indicated by a plus sign) and an output variable. Although edges are displayed unidirectional to represent the causality, messages will be passed in both directions.

### Implementation

EPIFANY starts with the output from probabilistic rescoring tools applied to peptide search engine results. It writes the PSM probabilities and associations from PSMs to proteins into OpenMS’ datastructures. After aggregating PSM probabilities on the peptide level and creating the bipartite graph from the peptides mapping to potential protein candidates as described in the previous section, the resulting protein-peptide graph is split into connected components by a depth-first search. Then—using the OpenMP 2.0 API^21^ in a dynamic scheduling mode—the processing of the connected components is distributed on as many CPU threads as available. For each connected component in the graph a factor graph is built, equivalent to the Bayesian network specified in the Model Section. The Bayesian network represented by the bipartite graph is converted to a factor graph as follows (example shown in Figure 2): Firstly, to allow querying posteriors for indistinguishable proteins, an additive factor is introduced for all proteins that share the same set of peptides. Then, to reduce loops, save computations, and avoid oscillations later on, we also create peptide cluster factors that hold the probabilities for the number of parent proteins for sets of peptides that share the same parent proteins or parent protein groups. Both of these factor types use convolution trees for an efficient adding of the probability tables. It is worth noting that this hierarchy could also include more intermediate factors but since no sub-quadratic algorithm is known to us to create an optimal hierarchy the algorithm only uses two levels of aggregation. Also, this aggregation is only performed if the number of contributing proteins or protein groups is greater than one. Finally, for each peptide a factor is added to the graph, that holds the CPD for the probability of a peptide being present given the number of parent proteins (see Model section, Equations 2 and 3). The factor nodes on the very left and very right are just singleton factors to keep track of the potentials (i.e., current "posteriors" during the LBP algorithm) on proteins and peptides. They are initialized with priors (for proteins) and evidence probabilities (for peptides) and after convergence of the algorithm hold the final posteriors to be queried. As mentioned earlier, by default the model parameters are evaluated based on a grid of possible values for each of them. While the rough skeleton of the factor graph is kept over multiple sets of parameters, its internals (e.g., the probability tables) are re-initialized for every combination. After performing message passing until convergence in a step-wise procedure—loosening convergence criteria as the number of messages passed increases—posteriors can be queried on all important levels: protein, protein group, or peptide level. All factors perform the same *p*-norm marginalization (see Equation 5) to generate a lower-dimensional message out of the higher-dimensional potentials. *p* can be freely chosen by the user. Even max-product inference (*p* = *∞*) is available at low additional computational costs^22^. A higher *p* is recommended together with the regularized model to have a positive effect on calibration when the degree of ambiguity in the database is suspected to be much higher than in the actual sample. Higher values of *p* result in messages that focus on the information from high-probability configurations (i.e., the proteins with the highest unique evidence).

An additional outer loop evaluates the model for points on a three-dimensional grid over the three parameters *α*, *β* and *γ* based on a convex combination of partial AUC (for target-decoy classification) and single protein posterior probability calibration (i.e., comparing posterior based and target-decoy based FDRs).

As an optional post-processing, a greedy protein group resolution can be performed. It implements a probabilistic maximum parsimony model, where protein groups are ordered by their posterior probability and starting from the best group, each greedily claims all peptides that it potentially generates until all peptides have been claimed. Proteins or protein groups without any remaining evidence are then deleted or implicitly assigned a probability of 0.

### Data and data pre-processing

To benchmark the main advancements of EPIFANY and to show the different strengths of the tool we focused on two different datasets.

Accuracy and calibration can best be measured on a dataset like the iPRG2016 benchmark data with a set of known ground truth Protein Epitope Signature Tags (PrESTs)^14^. Two sets (labelled A and B) of known PrESTs were designed to share a large number of peptides, spiked-into an *Escherichia coli* lysate background in three different experiments, then measured in triplicates: one experiment each exclusively containing one of the spike-in sets (A, B) and a third containing both sets of PrESTs (A+B). Our evaluation focuses on the task of identifying the PrESTs of (w.l.o.g.) B while having the full database of spike-ins (from A+B), entrapment proteins (i.e., intentionally absent PrESTs) and background proteins as potential candidates. All experiments contained an equimolar concentration of each PrEST. To avoid confusion, we will simply use the term “protein” instead of PrEST in the rest of the manuscript.

With the goal to measure scalability of the new tool a second, larger scale dataset was analyzed. It is part of an unpublished study and consists of two measurements of human cells on a long gradient at two different time points in duplicates.

#### iPRG2016 data set

Raw files from the corresponding PRIDE^23^ project PXD008425 were converted and centroided with msConvert^24^ on all levels. The `fasta` database provided together with the study was used to generate a decoy database through homology-aware (i.e., peptide-based) shuffling of amino acids with OpenMS’ DecoyDatabase^25,26^ tool. Then spectra were searched using Comet (2016.01 rev. 3) ^27^ allowing a 10 ppm precursor mass tolerance and one missed cleavage for fully tryptic peptides. As fixed modification we required Carbamidomethylation (C). Variable modification was set to Oxidation (M). After merging the results over replicates (by creating the union of proteins and concatenating PSMs) we added target-decoy annotations on protein- and PSM-level. To obtain better discrimination through Percolator 3.02 we extracted additional features specific for the Comet search-engine before running Percolator with standard settings for the PSM score re-calibration and basic protein inference activated. For all other methods tested we used the PSM-level posterior error probabilities reported by Percolator as input after filtering them slightly by removing PSMs with error probabilities higher or equal to 0.999 (to be consistent with the defaults in Fido and EPIFANY). For comparability with PIA and Percolator which only support group-level inference, Fido and EPIFANY were run with group-level inference as well, thereby querying posteriors on the (indistinguishable) protein group level (i.e., reporting the probability of at least one member being present). ProteinProphet and EPIFANY then report both, group-level and single protein-level probabilities.

#### Large-scale data

After searching the spectra of the four runs with MSGF+^28^ (unspecified number of missed cleavages, 10 ppm precursor mass tolerance, fully specific Trypsin/P as enzyme, fixed Methylthio (C) modification, variable Oxidation (M)) against the whole human part of the TrEMBL (incl. isoforms) database^29^ (with peptide-level pseudo-reversed decoys appended) and merging the samples the dataset can be summarized by the following numbers:

- 119,921 proteins
- 533,218 different peptide sequences
- 807,663 PSMs
- 734,522 spectra

Again, PSMs were re-scored and probabilities extracted via Percolator 3.02 by training its support vector machine on a subset of 250,000 PSMs for speed and memory efficiency. Since there is currently no way to run protein inference in Percolator separately, the corresponding options were set to be activated as well. To evaluate the computationally most demanding task for the Bayesian approaches, Fido was run in single protein level mode (no groups to exploit symmetries). EPIFANY calculates both levels simultaneously by default. Other parameters were left with their defaults in all tools, except for filtering out PSMs under 0.001 probability in ProteinProphet and PIA (to be comparable with the defaults in Fido and EPIFANY). Times and peak memory usage were measured with the Unix utility `time` on a two-socket Intel Xeon X5570 machine (i.e., 16 possible threads) with 64GB of RAM.

#### Benchmarked tools and tool-specific adaptions

The tools tested represent a diverse set of algorithms. PIA 1.3.10 was chosen as spectrum level parsimony approach considering shared peptides. Percolator 3.02 represents aggregation-based approaches on unique peptides only. ProteinProphet (compiled from TPP 5.1) was included as a commonly used iterative and pseudo-probabilistic heuristic which considers shared peptides as well. Lastly, Fido (in the version shipped with OpenMS) is the representative of Bayesian methods with a very similar model to EPIFANY’s but with a different inference procedure. The selection is also based on other recent evaluations of protein inference methods^14,30^.

Since PIA accepts OpenMS’ idXML format by default, the only change that was done was a renaming of the PSM score type name resulting from Percolator to OpenMS’ posterior error probability type so that PIA accepts the scores and interprets them as error probabilities.

For ProteinProphet OpenMS’ IDFileConverter was used to write the error probabilities from Percolator into a pepXML file that is compatible with ProteinProphet to make sure that all tools start from the same set of scores. The FDR estimation procedures of the tools were used whenever possible. For ProteinProphet we used the same FDR estimation procedure as for EPIFANY with a concatenated target-decoy database and the equation 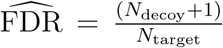^31^. Groups are counted as decoy if they consist of only decoy proteins. Reported are always q-values unless stated otherwise.

## Results and discussion

### EPIFANY shows improved identification performance

Results on the unpooled experiments of the iPRG2016 study show that EPIFANY (with greedy group resolution enabled) yields the highest count of known true positive proteins among the tested methods (Figure 3). On this dataset EPIFANY reaches a 9.78% higher true positive count at 5% FDR than the second best method Percolator. Considering the few missing true positives to be identified, this increase is of even greater importance. PIA and ProteinProphet did not perform well on this barely filtered set of PSMs with ProteinProphet’s reported proteins all having q-values higher than 5%. The performance of ProteinProphet might be explained by the different pipeline^32^ usually used to create its input. Furthermore, ProteinProphet performs a different and more aggressive protein grouping by not only aggregating indistinguishable groups but also by subsuming groups into more general protein ambiguity groups.

**Figure 3:**
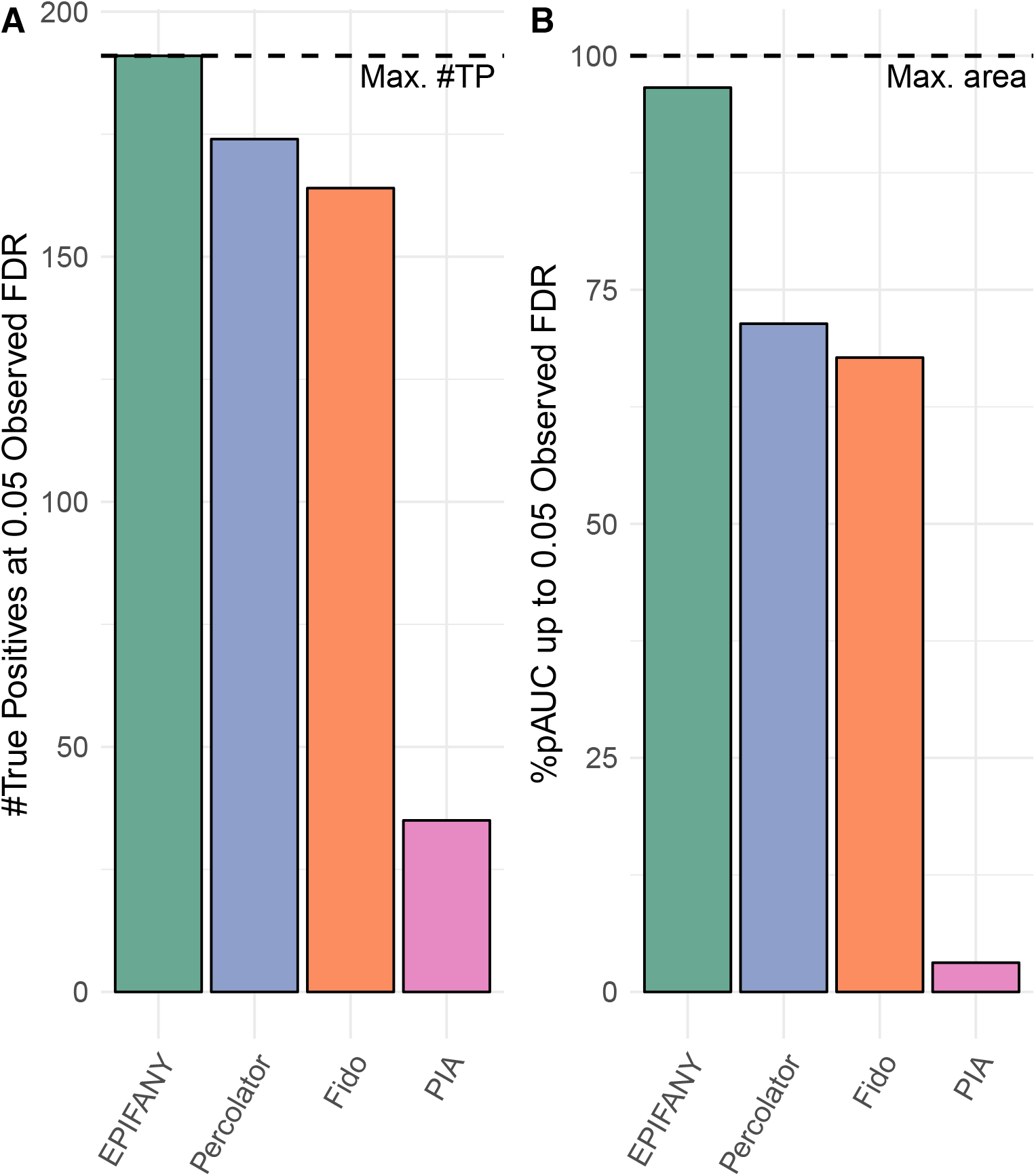
Summary statistics on the identification performance of different inference methods at a 5% entrapment FDR on the iPRG2016 dataset, sample “B”. ProteinProphet was not included in this comparison as it yielded first true identifications at an FDR above 5% only. **A)** Number of true positive proteins found. The maximum number of true positives according to the ground-truth database given is 191 and indicated by a dashed horizontal line. **B)** The percentage of the maximum area under the partial receiver operating curve (%pAUC) as a measure of how quickly methods accumulate true positives at increasing FDR levels until the chosen cutoff at 5% FDR.

However, since this dataset is limited in the number of known proteins that can be found (191 in sample B), looking at the number of identifications at a certain cutoff does not show the full strength of our new method. Not only does it identify more known spike-ins, it also finds them at overall lower FDRs than every other method, and thus at a threshold where it is reporting fewer false positives. This is evident in the quickly rising receiver operating curve (ROC), covering the largest partial area under the curve (truncated at 5% FDR; abbreviated pAUC) of all tools. This is summarized in Figure 3B. Together with Fido it is also the only tool reporting all correct proteins after all—though Fido finds all 191 present proteins at an FDR of 47% (beyond the cutoff in the figures). Other tools (even without any FDR cutoff) completely ignore or filter some true positives, most likely due to missing/insufficient evidence, e.g., missing unique peptides in Percolator (which reaches 187 true proteins at its maximum FDR of 56%).

When applied to data without stringent PSM FDR filtering, some tools (e.g., PIA) achieved only poor pAUCs. Therefore we performed the evaluation for all tools again with a suggested pre-cutoff of 0.01 PSM FDR^33^. Most methods benefit from pre-filtering in various degrees. PIA now performs almost identically to EPIFANY (Supplementary Figure S2). While a filtering on PSM-level before inference indeed seems beneficial to identification performance on this dataset we hypothesize that on other datasets with noisier data (e.g., sub-optimal or overly permissive search engine settings with many considered modifications) this cutoff is too conservative and in the end one is missing out on correct identifications with low-scoring evidence under this cutoff. Good performance on both strictly filtered and almost unfiltered data therefore shows the robustness of our method.

### FDR estimates of EPIFANY are more realistic than other estimates

Before introduction of a regularized model and greedy resolution as implemented in EPIFANY, Bayesian methods were shown to report overly optimistic FDRs in the case of datasets like the one tested here^14^. This is due to the fact that although only a small number of proteins are known to be present in the sample the spectra were searched against an additional equal number of absent proteins designed to share peptides with the present ones plus an even bigger number of 1,000 absent random entrapment proteins. Unregularized models using standard sum-product inference like Fido then would conservatively assign probabilities far from 0 to proteins with similar or no unique evidence which share one or more peptides with truly present proteins. Regularized max-product inference in EPIFANY now makes the assumption that it is less likely for peptides to be generated by many proteins and preferentially distributes the evidence among proteins with the highest unique evidence. Greedy resolution additionally makes a definite, unprobabilistic choice among those proteins based on their posterior probability. By postponing this decision until the end of an identification pipeline, however, reliable uncertainty estimates are available up to the very last step. Compared to methods ignoring shared peptides for FDR estimation (e.g., Percolator), FDRs reported by EPIFANY are closer to the true FDRs. The differences between reported and observed FDR can be seen in Figure 4 which shows them up to a 15% FDR cutoff (since higher cutoffs are usually not of interest). The cutoff was set this high to generate more robust measures of calibration (by being able to include more point estimates for each method). Additional ROCs and calibration plots on the full FDR range can be found in Supplementary Figure S1.

**Figure 4:**
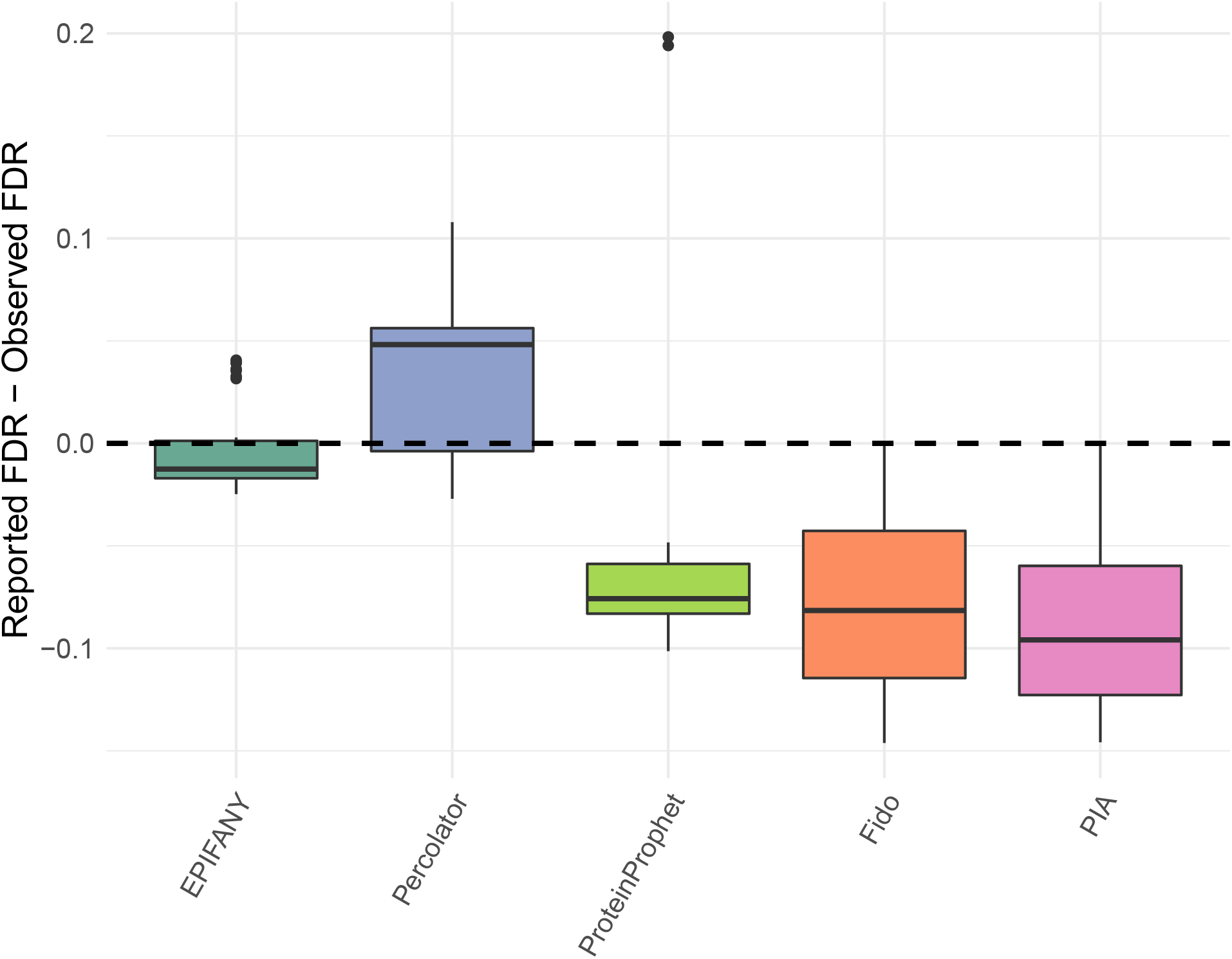
Absolute deviations from reported FDR from different tools and settings to the true entrapment FDR (on the subset of proteins with known presence/absence) on the iPRG2016 dataset (experiment “B”). The boxplots aggregate data points for each method until a 15% entrapment FDR. Lower deviations imply better calibration with perfect calibration being indicated by the dashed horizontal line. Data above the horizontal line signify conservative estimates while data below the line signify overly optimistic estimates.

### Scalable algorithms allow application to large-scale datasets with vastly disparate discoveries

A common challenge with inference on generative Bayesian models was their scalability due to the speed of the method, which is inherently tied to the complexity of the underlying model. The efficiency of EPIFANY enables full Bayesian inference on problem sizes that were previously intractable given the used model. On the human dataset searched based on the TrEMBL database with isoforms even non-Bayesian approaches struggle with the high connectivity in the resulting protein-peptide graph. This is evident in Figure 5 which shows the runtimes and memory consumption of the different tools. Fido took longer than a day of runtime, ProteinProphet did not finish after a week and also PIA required more than two days of processing (without considering compilation of the intermediate graph format used by PIA). While multi-threaded EPIFANY (16 min) is even faster than (subset-trained) Percolator (33 min), it should be noted that most of Percolator’s runtime consists of peptide rescoring and internal training of a support vector machine model. The same argument holds for memory usage. Although running Percolator just for protein inference is to our knowledge currently not possible it is likely that the actual runtime on that dataset will be in the single-digit minutes.

**Figure 5:**
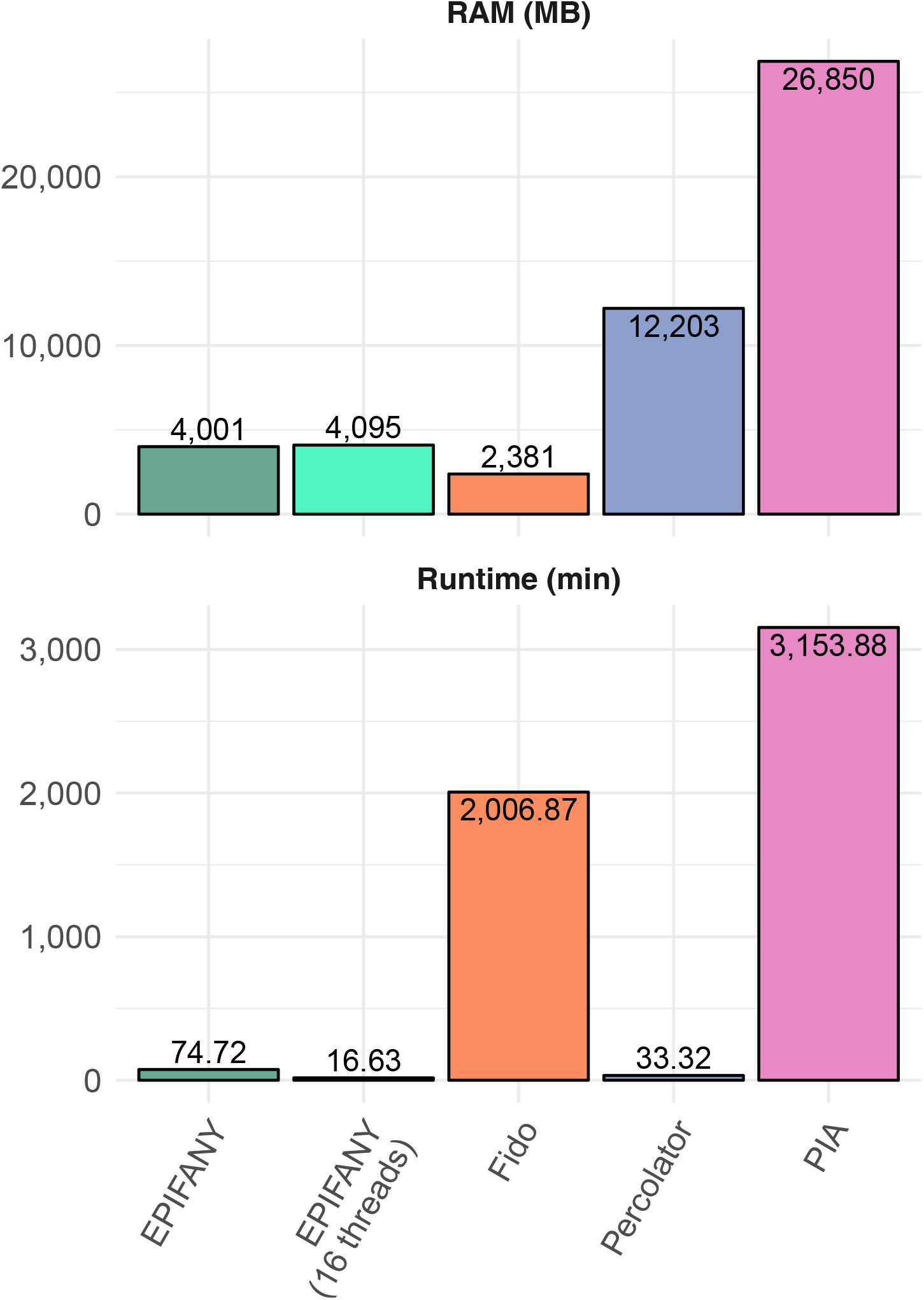
Memory usage in megabytes (upper) and runtimes in minutes (lower) on the large-scale human dataset filtered at 0.001 PSM probability. ProteinProphet did not terminate within a week of runtime and was thus not included in the figure.

We also emphasize, that an application of a Bayesian method incorporating results from shared peptides may lead to different discoveries compared to methods currently used on big datasets that ignore this complication^7,34^. On the larger of the tested datasets for example, Percolator considers only half of the roughly 50,000 potential target proteins (with at least one PSM above the 0.001 probability cutoff as used in EPIFANY) due to missing unique evidence for the rest of them (data not shown).

### Implementation grants flexibility in input and output information

In addition to increased performance, EPIFANY’s general Bayesian framework also permits the inclusion of auxiliary information (besides PSM data) in a convenient manner. The new implementation allows the usage of arbitrary protein priors for any protein which allows integration of information from complementary RNA-seq experiments, which has been shown to improve protein identification^35^. Due to its modular graph structure it is also easy to add additional probabilistic evidence on the peptide level in the future. Since the factor graph includes both single proteins and indistinguishable protein groups, both results can be output simultaneously. Single protein-level probabilities of proteins in indistinguishable groups are e.g., not possible to be reported in PIA or Percolator. Another novelty is the reporting of updated peptide-level posteriors that can be used to re-score and improve FDR on peptide-level, yielding increased target-decoy classification performance by up to 6% (in the area under the receiver operating curve). This effect comes from the fact that the information from sibling peptides^9^ in the graph was now propagated and incorporated into every peptide posterior.

### Limitations

Since the algorithm’s grid search explores multiple parametrizations of the model, there might be settings in which at some point during LBP contradicting messages are generated that have disagreeing beliefs about the probability of a protein or peptide (e.g., from evidence of different parts of the graph). This can often be solved by a step-wise increase in dampening in later iterations. In extreme cases, however, it can lead to interruptions in the inference on that part of the graph (since for example zero probabilities cannot be recovered). In the latter case we evaluate the (failing) parameter set by using the prior probabilities of the affected proteins (assuming no knowledge could be gained for those proteins). In the case of non-convergence due to too many iterations the current beliefs for a protein are used. However, those cases are rare and usually seen in very large components caused by extreme parameter sets only.

Also, due to the fact that the parameter estimation is based on target-decoy annotations, our method is affected by the composition of the decoy database. However, as shown in Supplementary Figure S3 different random shuffles of the iPRG2016 database resulted in comparable identification performances with a median partial AUC of 96% and a median absolute deviation of one percent point.

Furthermore, group-level inference is still based on *experimentally* indistinguishable proteins which hinders reproducibility of the specific groupings across multiple runs.^6^ If other types of groupings should be performed this has to be reflected in the input before running inference (e.g., by merging protein IDs beforehand).

### Availability

EPIFANY runs on all major platforms (Windows, Linux, OSX). It is available under an open-source license (BSD three-clause) at https://openms.org/epifany. This website also contains demo data, a manual, binary installers for all platforms and links to the source code repository. EPIFANY relies on the Evergreen inference library released by O. Serang under an MIT license which can be found under https://bitbucket.org/orserang/evergreenforest^36,37^

## Conclusion and Outlook

With EPIFANY we present a new approach to efficient Bayesian protein inference in proteomics that combines excellent inference quality with good runtimes. The underlying method certainly can be improved upon. As mentioned in the Limitations Section, the results depend on the convergence of the algorithm. Convergence is generally affected by the current parametrization of the model as well as the connectivity in the graph. To which degree still has to be investigated. In case of sub-optimal results on a connected component the algorithm could in the future try different message scheduling types or resort to heuristics. Additionally, the currently experimental options of peptide re-scoring and user-defined priors are worth further research. Incorporation of additional evidence especially from MS1-level, replicates or multiple PSMs is a viable extension, too, however, initial tests with precursor mass and retention time deviations yielded noisy, generally disappointing results so far. Regularization of protein groups could be improved by facilitating user-defined protein groups (e.g., by gene or theoretical digest). Together with a re-introduction of proteotypicity with discretized *α* parameters per peptide instead of a single *α* per dataset it could help in the discrimination of otherwise indistinguishable proteins. In general, parameter estimation via grid-search is a very time-consuming part of the algorithm and although different parameter sets can be distributed across machines on even larger-scale data, learning speed might be improved by learning on a subset of the graph or completely circumvented by including the parameters as hyperparameters into the probabilistic model.

## Acknowledgement

First of all the authors thank Xiao Liang for the fruitful discussions and ideas on this and related topics as well as implementations in earlier stages of this work and for the happy atmosphere she brought to the office. The authors also thank Julian Uszkoreit for the generation of the input graph file on the large dataset for his tool PIA and his help with the settings.

Furthermore the authors are thankful for the support of the funding agencies and their following grants:

- J.P., T.S. and O.K. were funded by the German Federal Ministry of Education and Research (BMBF) under FKZ 031A535A (German Network for Bioinformatics).
- O.S. received funding from the University of Montana small grant 325476 (UGP 2018: Development of Probabilistic Cardinal Models) and the National Institute of General Medical Sciences of the National Institutes of Health under Award Number P20GM103546. This material is based upon work supported by the National Science Foundation under grant no. 1845465.
- T.D. was funded by the European Union through the EU-H2020 ICT-644727 COGIMON project.
- Funding for O.K. was also provided by EU project EPIC-XS grant number 823839 (H2020-INFRAIA)

## Supporting Information Available

- SupplementaryFigures.docx: Supplementary Figures S1-3 comprising more detailed information on the iPRG2016 results (Suppl. Fig. S1), plus implications of prefiltering PSMs (Suppl. Fig. S2) as well as the effect of differently shuffled decoy databases on EPIFANY’s parameter estimation (Suppl. Fig. S3). Supplementary figures S1 and S2 also include results of different EPIFANY settings.

## Author Contributions

O.S., K.R. and O.K. designed the experiments. O.S. and J.P. wrote the implementation of the method. J.P. performed data analysis and wrote the manuscript. T.S. wrote auxiliary tools and functions for data preprocessing and export. J.P., O.S., O.K. and T.D. interpreted the results. All authors helped in drafting and read as well as approved the final manuscript.

## References

(1) Zhang, Y.; Fonslow, B. R.; Shan, B.; Baek, M.-C.; Yates, J. R. Protein Analysis by Shotgun/Bottom-up Proteomics. Chemical Reviews 2013, 113, 2343–2394.

(2) Nesvizhskii, A. I.; Aebersold, R. Interpretation of shotgun proteomic data: the protein inference problem. Molecular & cellular proteomics: MCP 2005, 4, 1419–40.

(3) Serang, O.; Noble, W. A review of statistical methods for protein identification using tandem mass spectrometry. Statistics and its interface 2012, 5, 3–20.

(4) Uszkoreit, J.; Maerkens, A.; Perez-Riverol, Y.; Meyer, H. E.; Marcus, K.; Stephan, C.; Kohlbacher, O.; Eisenacher, M. PIA: An Intuitive Protein Inference Engine with a Web-Based User Interface. Journal of Proteome Research 2015, 14, 2988–2997.

(5) Cottrell, J. S. Protein identification using MS/MS data. Journal of Proteomics 2011, 74, 1842–1851.

(6) Serang, O.; Moruz, L.; Hoopmann, M. R.; Käll, L. Recognizing uncertainty increases robustness and reproducibility of mass spectrometry-based protein inferences. Journal of proteome research 2012, 11, 5586–91.

(7) The, M.; MacCoss, M. J.; Noble, W. S.; Käll, L. Fast and Accurate Protein False Discovery Rates on Large-Scale Proteomics Data Sets with Percolator 3.0. Journal of The American Society for Mass Spectrometry 2016, 27, 1719–1727.

(8) Käll, L.; Canterbury, J. D.; Weston, J.; Noble, W. S.; MacCoss, M. J. Semi-supervised learning for peptide identification from shotgun proteomics datasets. Nature Methods 2007, 4, 923–925.

(9) Nesvizhskii, A. I.; Keller, A.; Kolker, E.; Aebersold, R. A Statistical Model for Identifying Proteins by Tandem Mass Spectrometry. Analytical Chemistry 2003, 75, 4646–4658.

(10) Serang, O.; MacCoss, M. J.; Noble, W. S. Efficient marginalization to compute protein posterior probabilities from shotgun mass spectrometry data. Journal of proteome research 2010, 9, 5346–57.

(11) Li, Y. F.; Arnold, R. J.; Li, Y.; Radivojac, P.; Sheng, Q.; Tang, H. A bayesian approach to protein inference problem in shotgun proteomics. Journal of computational biology: a journal of computational molecular cell biology 2009, 16, 1183–93.

(12) Gerster, S.; Qeli, E.; Ahrens, C. H.; Bühlmann, P. Protein and gene model inference based on statistical modeling in k-partite graphs. Proceedings of the National Academy of Sciences of the United States of America 2010, 107, 12101–6.

(13) Pearl, J. Probabilistic reasoning in intelligent systems: networks of plausible inference; Morgan Kaufmann Publishers Inc.: San Mateo, CA, 1988.

(14) The, M.; Edfors, F.; Perez-Riverol, Y.; Payne, S. H.; Hoopmann, M. R.; Palmblad, M.; Forsström, B.; Käll, L. A Protein Standard That Emulates Homology for the Characterization of Protein Inference Algorithms. Journal of Proteome Research 2018, 17, 1879–1886.

(15) Berrou, C.; Glavieux, A.; Thitimajshima, P. Near Shannon limit error-correcting coding and decoding: Turbo-codes. 1. Proceedings of ICC ’93 – IEEE International Conference on Communications. 1993; pp 1064–1070.

(16) Murphy, K. P.; Weiss, Y.; Jordan, M. I. Loopy Belief Propagation for Approximate Inference: An Empirical Study. Proceedings of the Fifteenth Conference on Uncertainty in Artificial Intelligence. San Francisco, CA, USA, 1999; pp 467–475.

(17) Pearl, J. Reverend Bayes on Inference Engines: A Distributed Hierarchical Approach. Proceedings of the National Conference on Artificial Intelligence, Pittsburgh, PA, USA, August 18–20, 1982. 1982; pp 133–136.

(18) Serang, O. The probabilistic convolution tree: efficient exact Bayesian inference for faster LC-MS/MS protein inference. PloS one 2014, 9, e91507.

(19) Elidan, G.; McGraw, I.; Koller, D. Residual Belief Propagation: Informed Scheduling for Asynchronous Message Passing. Proceedings of the Twenty-Second Conference on Uncertainty in Artificial Intelligence 2006. 2006.

(20) Jensen,; Lauritzen, S.; Olesen, K. Bayesian updating in causal probabilistic networks by local computations. Computational Statistics Quaterly 1990, 4, 269–282.

(21) OpenMP Architecture Review Board, OpenMP Application Program Interface Version 2.0. 2002; https://www.openmp.org/wp-content/uploads/cspec20.pdf.

(22) Pfeuffer, J.; Serang, O. A Bounded p-norm Approximation of Max-Convolution for Sub-Quadratic Bayesian Inference on Additive Factors. Journal of Machine Learning Research 2016, 17, 1–39.

(23) Perez-Riverol, Y. et al. The PRIDE database and related tools and resources in 2019: improving support for quantification data. Nucleic acids research 2019, 47, D442–D450.

(24) Chambers, M. C. et al. A cross-platform toolkit for mass spectrometry and proteomics. Nature Biotechnology 2012, 30, 918–920.

(25) Sturm, M.; Bertsch, A.; Gröpl, C.; Hildebrandt, A.; Hussong, R.; Lange, E.; Pfeifer, N.; Schulz-Trieglaff, O.; Zerck, A.; Reinert, K.; Kohlbacher, O. OpenMS âĂŞ An open-source software framework for mass spectrometry. BMC Bioinformatics 2008, 9, 163.

(26) Röst, H. et al. OpenMS: A flexible open-source software platform for mass spectrometry data analysis. Nature Methods 2016, 13, 741–748.

(27) Eng, J. K.; Jahan, T. A.; Hoopmann, M. R. Comet: An open-source MS/MS sequence database search tool. PROTEOMICS 2013, 13, 22–24.

(28) Kim, S.; Gupta, N.; Pevzner, P. A. Spectral probabilities and generating functions of tandem mass spectra: a strike against decoy databases. Journal of proteome research 2008, 7, 3354–63.

(29) Bairoch, A.; Apweiler, R. The SWISS-PROT protein sequence database and its supplement TrEMBL in 2000. Nucleic acids research 2000, 28, 45–48.

(30) Audain, E.; Uszkoreit, J.; Sachsenberg, T.; Pfeuffer, J.; Liang, X.; Hermjakob, H.; Sanchez, A.; Eisenacher, M.; Reinert, K.; Tabb, D. L.; Kohlbacher, O.; Perez-Riverol, Y. In-depth analysis of protein inference algorithms using multiple search engines and well-defined metrics. Journal of Proteomics 2017, 150, 170–182.

(31) Levitsky, L. I.; Ivanov, M. V.; Lobas, A. A.; Gorshkov, M. V. Unbiased False Discovery Rate Estimation for Shotgun Proteomics Based on the Target-Decoy Approach. Journal of Proteome Research 2017, 16, 393–397.

(32) Deutsch, E. W.; Mendoza, L.; Shteynberg, D.; Slagel, J.; Sun, Z.; Moritz, R. L. Trans-Proteomic Pipeline, a standardized data processing pipeline for large-scale reproducible proteomics informatics. PROTEOMICS – Clinical Applications 2015, 9, 745–754.

(33) Uszkoreit, J.; Perez-Riverol, Y.; Eggers, B.; Marcus, K.; Eisenacher, M. Protein Inference Using PIA Workflows and PSI Standard File Formats. Journal of Proteome Research 2019, 18, 741–747.

(34) Savitski, M. M.; Wilhelm, M.; Hahne, H.; Kuster, B.; Bantscheff, M. A Scalable Approach for Protein False Discovery Rate Estimation in Large Proteomic Data Sets. Molecular & Cellular Proteomics 2015, 14, 2394–2404.

(35) Proffitt, J. M.; Glenn, J.; Cesnik, A. J.; Jadhav, A.; Shortreed, M. R.; Smith, L. M.; Kavanagh, K.; Cox, L. A.; Olivier, M. Proteomics in non-human primates: utilizing RNA-Seq data to improve protein identification by mass spectrometry in vervet monkeys. BMC Genomics 2017, 18, 877.

(36) Serang, O. The p-convolution forest: a method for solving graphical models with additive probabilistic equations. arXiv e-prints 2017, arXiv:1708.06448.

(37) Lucke, K.; Thibeau, M.; Pfeuffer, J.; Liang, X.; Serang, O. The Titin Problem: Hitch-hiking Siblings and an Engine for Experimenting with Protein Inference Models. 2019; (in preparation).

