## Supplementary Figures 1-3 for "EPIFANY – A method for efficient high-confidence protein inference"

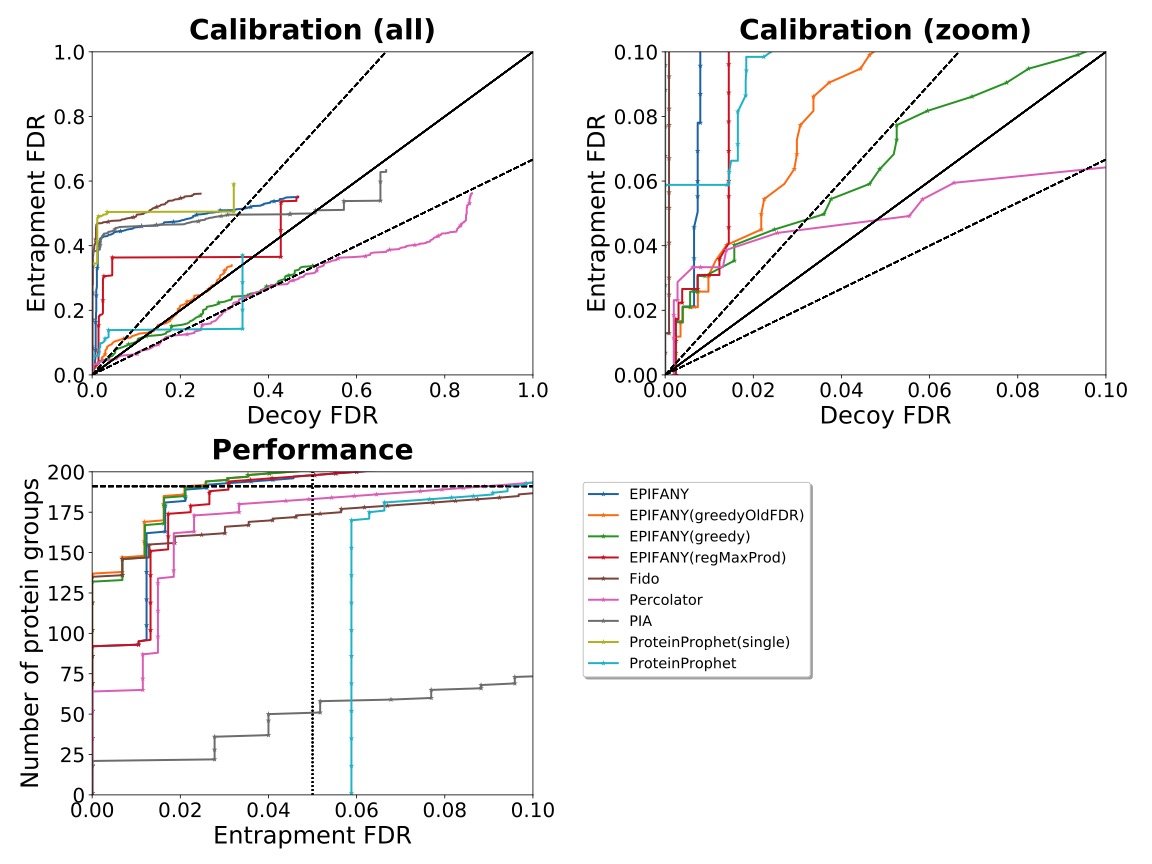

Supplementary Figure S1: Results on the “B” sample from the official iPRG2016 Evaluation script (adapted to show multiple methods). The log-log calibration subfigure was removed in favor of a legend. Combined PSMs of the three replicates were filtered to include only PSMs with >=0.001 Percolator probability. “RegMaxProd” means a regularized model with max product inference was used. “Greedy” in the label shows that greedy group resolution based on posteriors was enabled. Methods with “OldFDR” use a different equation to estimate FDRs based on target-decoy estimation (D/(T+D) instead of the more conservative (D+1)/T).

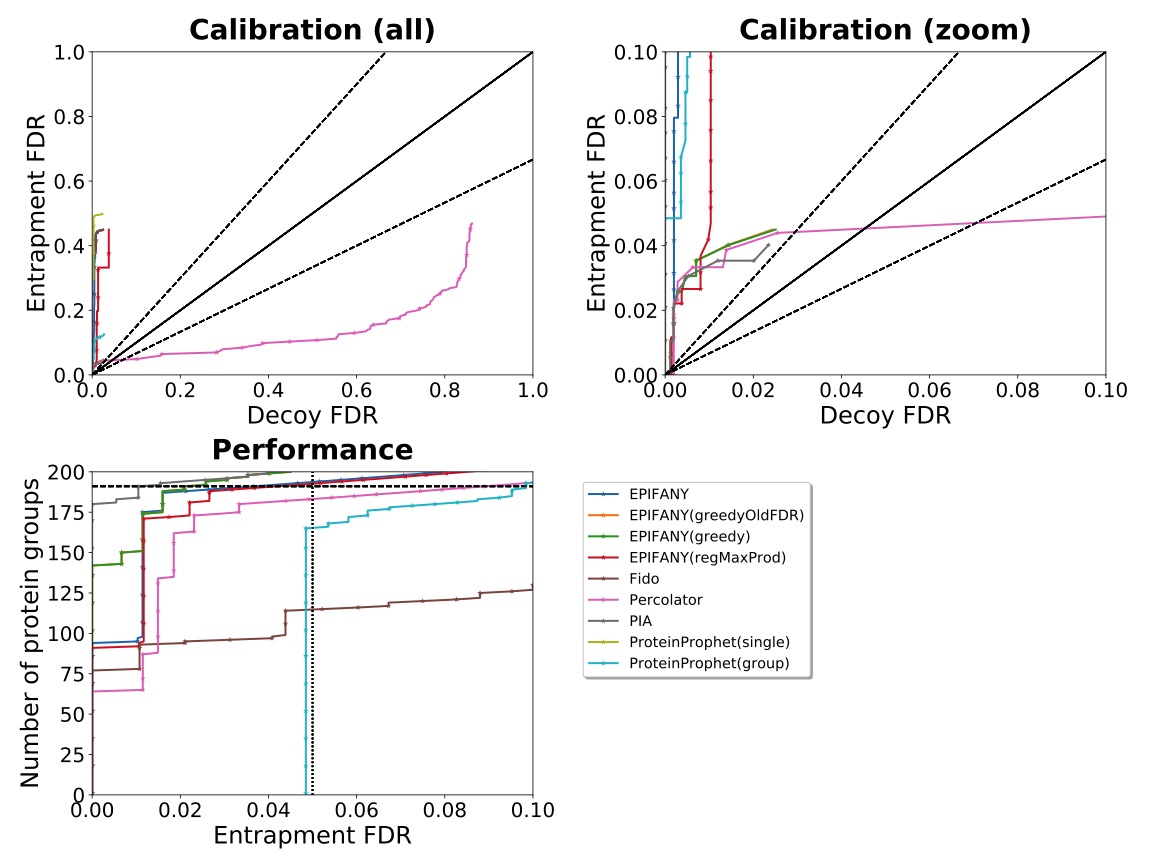

Supplementary Figure S2: Results on the “B” sample from the official iPRG2016 Evaluation script (adapted to show multiple methods). The log-log calibration subfigure was removed in favor of a legend. Combined PSMs of the three replicates were filtered for 1% (concatenated) target-decoy FDR on PSM level. “RegMaxProd” means a regularized model with max product inference was used. “Greedy” in the label shows that greedy group resolution based on posteriors was enabled. Methods with “OldFDR” use a different equation to estimate FDRs based on target-decoy estimation (D/(T+D) instead of the more conservative (D+1)/T).

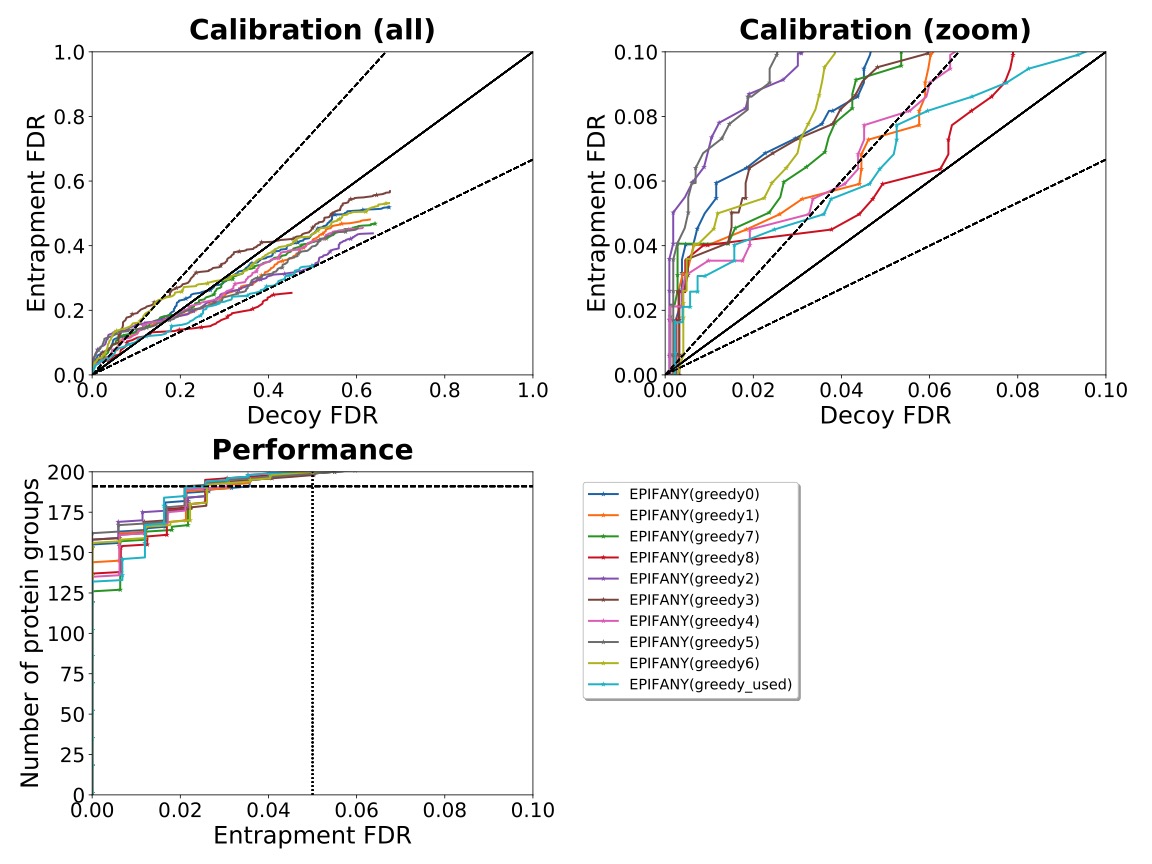

Supplementary Figure S3: Results on the “B” sample from the official iPRG2016 Evaluation script (adapted to show multiple methods). The log-log calibration subfigure was removed in favor of a legend. Combined PSMs of the three replicates were filtered to include only PSMs with >=0.001 Percolator probability. Different runs of the Comet, Percolator, EPIFANY pipeline were carried out with differently seeded, randomly shuffled decoy databases to show the robustness of the parameter estimation based on protein target-decoy FDRs. Different best parameters were estimated in the repeated runs but spanned a narrow subspace of the considered grid. Larger difference in the calibration can be explained by the relatively large impacts of the ranks of false positives on such a small set of true positives. The randomly chosen run labeled “greedy_used” used the same decoy database as in Supplementary Figures 1 and 2 and is evaluated in the manuscript.
